# Metabolic biomarker-based phenotyping unveils quantitative effects of plant resistance and pathogen aggressiveness in the grapevine (Vitis spp.) - downy mildew (Plasmopara viticola) pathosystem

**DOI:** 10.1101/2025.09.25.678557

**Authors:** Tyrone Possamai, Raymonde Baltenweck, Sabine Wiedemann-Merdinoglu, Marie-Céline Lacombe, Marie-Annick Dorne, Matéo Bareyre, Erik Griem, René Fuchs, Jochen Bogs, Éric Duchêne, Pere Mestre, Didier Merdinoglu, Philippe Hugueney

**Author notes:** Corresponding authors (TP); (RB).

## Abstract

Grapevine resistance to downy mildew has been primarily associated with major “Resistance to *Plasmopara viticola*” (*Rpv*) loci, which are extensively used in breeding programs. Resistant varieties represent an effective solution to mitigate the environmental impact of fungicide application in viticulture, but *P. viticola* strains able to overcome major *Rpv* have become a main threat to their cultivation. Pyramiding resistance loci in the same variety enhances plant resistance, but interactions involving stacked and defeated *Rpv* and different *P. viticola* strains are poorly documented. Investigation of these interactions may uncover new information for the development of efficient breeding strategies, the optimal exploitation of *Rpv,* and the building of durable resistance. In the present study, a grapevine offspring carrying single and pyramided *Rpv1*, *Rpv3.1* and *Rpv10* was phenotyped in laboratory conditions for the resistance to *P. viticola* using a naive strain and a strain virulent towards *Rpv10*. By using a high-resolution phenotyping strategy based on *P. viticola* metabolic biomarkers, we demonstrated that the efficacy of *Rpv* combinations and aggressiveness of *P. viticola* strains can be quantified in the early phase of infection and were often related to sporulation outcome. Furthermore, we described how a limited residual effect of a defeated *Rpv* may become significant in pyramiding. In conclusion, in addition to providing the keys to streamlining resistance utilization in grapevine, our research presents a distinctive case study that provides valuables information for breeding new resistant varieties, thanks to an innovative “omic”-based phenotyping approach, which may be adapted to other plant pathosystems.

## Introduction

Grapevine (*Vitis* spp.) is a fruit crop cultivated worldwide for table grape, wine and raisin production [1]. The oomycete *Plasmopara viticola* is the causal agent of grape downy mildew. Its control is usually based on the recurrent use of agrochemicals, which have adverse human health, socio-economic and environmental impacts [2]. *P. viticola* has shown the capacity to develop resistance to a wide range of fungicides and represents a significant threat for viticulture [3]. The breeding of new cultivars resistant to pathogens is an effective solution to reduce the use of agrochemicals, control plant diseases and enhance agriculture sustainability [4,5]. Over the past two decades, more than 30 grapevine resistance loci to *P. viticola* (*Rpv*) [6] have been identified and some major loci, like *Rpv1*, *Rpv3*, *Rpv10* and *Rpv12*, have been successfully introgressed into cultivated grapevines and represent common genetic resources implemented in breeding [5,7]. Major *Rpv*-mediated resistance has been associated to nucleotide-binding leucine-rich repeat (NLR) genes [8,9], a main type of resistance genes involved in effector-triggered immunity (ETI) [10]. According to the gene-for-gene concept, the efficacy of major genes can be affected by the specific combination of virulence genes found in different pathogen strains [10]. The pathogen *P. viticola* has demonstrated a high evolutionary potential, with strains isolated from resistant hosts exhibiting a greater aggressiveness, characterized by a short latency period and high levels of spore production [11]. Furthermore, virulent strains that completely breakdown one or more *Rpv* have been identified and now pose a significant threat to the cultivation of new resistant varieties [12,13]. Combining several resistance genes (pyramiding) in the same variety is an established strategy to enhance the efficacy, stability and durability of plant resistances [14,15], and a correlation between the resistance level and the number of resistance genes has been observed in grapevine [16].

The phenotyping of grapevine-downy mildew interaction is usually based on the visual evaluation of *P. viticola* sporulation, which emerges on the abaxial surface of the leaf, and of grapevine cells necrosis [7,12], which are indicative of plant’s recognition of the pathogen and deployment of an active plant response [17]. However, visible symptoms are often assessed by human experts using categorical scales, whose limited resolution and potential subjectivity complicate the interpretation and comparison of results. [7]. Image analysis represents a quantitative and objective phenotyping method, but it can be affected by interferences from water droplets, leaf morphology and hairiness as well as poorly resolved pathogen structures [7]. Finally, phenotyping methods based on visible symptoms are only applicable in advanced stages of infection. Recently, *P. viticola*-specific metabolites have been identified as biomarkers for the monitoring and quantification of the pathogen in grapevine leaves [18]. Metabolic biomarkers represent a new tool to overcome some issues in grapevine resistance and *P. viticola* interactions phenotyping, towards a better quantification of plant resistance and pathogen aggressiveness.

In this study, the potential of single and pyramided combinations of *Rpv1*, *Rpv3.1* and *Rpv10* to enhance grapevine resistance against *P. viticola* was investigated. Plants carrying all possible combinations of these *Rpvs* were challenged with a naive pathogen strain, and a strain virulent on *Rpv10*, termed *avrRpv10-*. The outcomes of the interactions between the *Rpv* combinations and *P. viticola* strains were characterized by integrating the visual assessment with the high-resolution quantification of *P. viticola*-specific metabolic biomarkers. This comprehensive approach yields new information on the effectiveness of single and pyramided *Rpv* and provides a better understanding of *P. viticola* strains’ adaptation to the host, with a view toward the construction of effective and durable resistances in grapevine breeding programs. Moreover, this work validates the use of metabolomics-based phenotyping for the precise characterization of quantitative aspects of plant genetic resistance and pathogen aggressiveness.

## Results

### Visual assessment of *P. viticola* infection

A total of eight *Rpv* combinations based on *Rpv1*, *Rpv3.1* and *Rpv10* were tested in laboratory conditions (3 genotypes/*Rpv* combination), under parallel inoculations with a naive *P. viticola* strain and a strain virulent towards *Rpv10* (16 *Rpv*-*P. viticola* interactions). The inoculated leaf discs (3 discs/genotype/strain) were visually assessed between 3 and 6 days-post inoculation (dpi) for the incidence and severity of *P. viticola* sporulation by using the variable OIV452-1 [19,20], and for the incidence of necrosis (overview of the study and experimental design in Supplementary File S1).

The progression of the *P. viticola* infection and the outcomes at 6 dpi were found to depend on the *Rpv* combination and strain under consideration (Fig. 1; Table 1; Supplementary File S2) in two experiment replicates (Supplementary File S3-4a). A different aggressiveness of the *P. viticola* naive and *avrRpv10-* strains was observed on all the studied *Rpv* combinations (Fig. 2; Supplementary File 4b). Thus, the *Rpv*-*P. viticola* strain interactions for the two strains were separately analysed.

**Figure 1.**
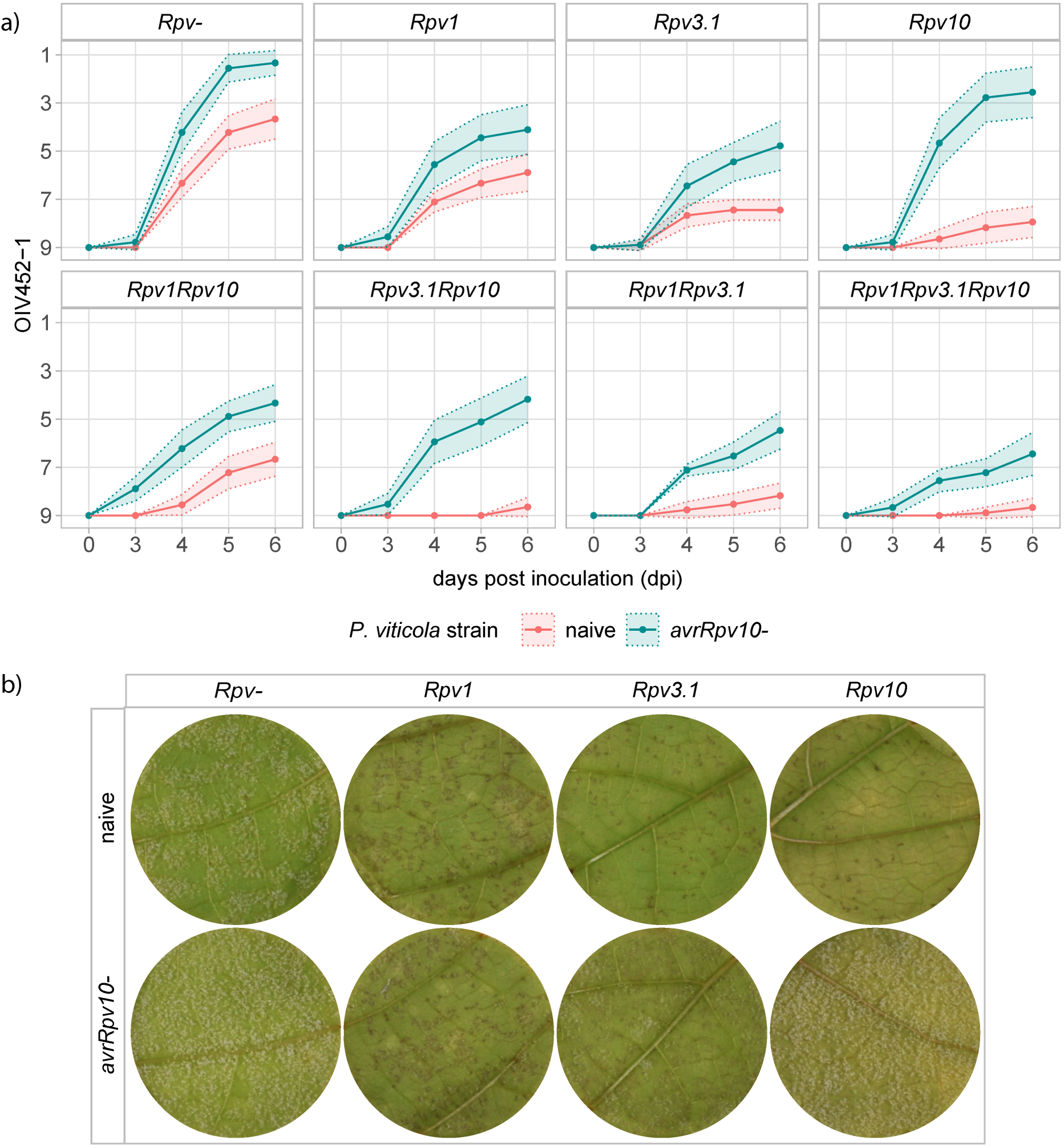
**Grapevine leaf discs infected with *P. vitico/a.*** a) OIV452-1 mean scores (points) and confidence limits (dashed area; 95%) describing the severity of *P vitico/a* infection on leaf discs (1 = sporulation densely covering the whole disc; 9 = absence of sporulation), collected for the different *Rpv* combinations under infection with the two *P vitico/a* strains. Data for 3 genotypes/Rpv combina­ tion and 3 discs /genotype/strain/dpi of two experiments are displayed. b) Representative discs at 6 dpi for the non-resistant *(Rpv-)* and single *Rpv* combinations inoculated with the two *P vitico/a* strains evidencing different rates of pathogen sporulation and grapevine necrotic response depen­ ding on the *Rpv-P viticola* interaction. OIV452-1 mean scores for single experiments are available in Supplementary File S3, and images of the discs at 3-4-5-6 dpi in Supplementary File S2.

**Figure 2.**
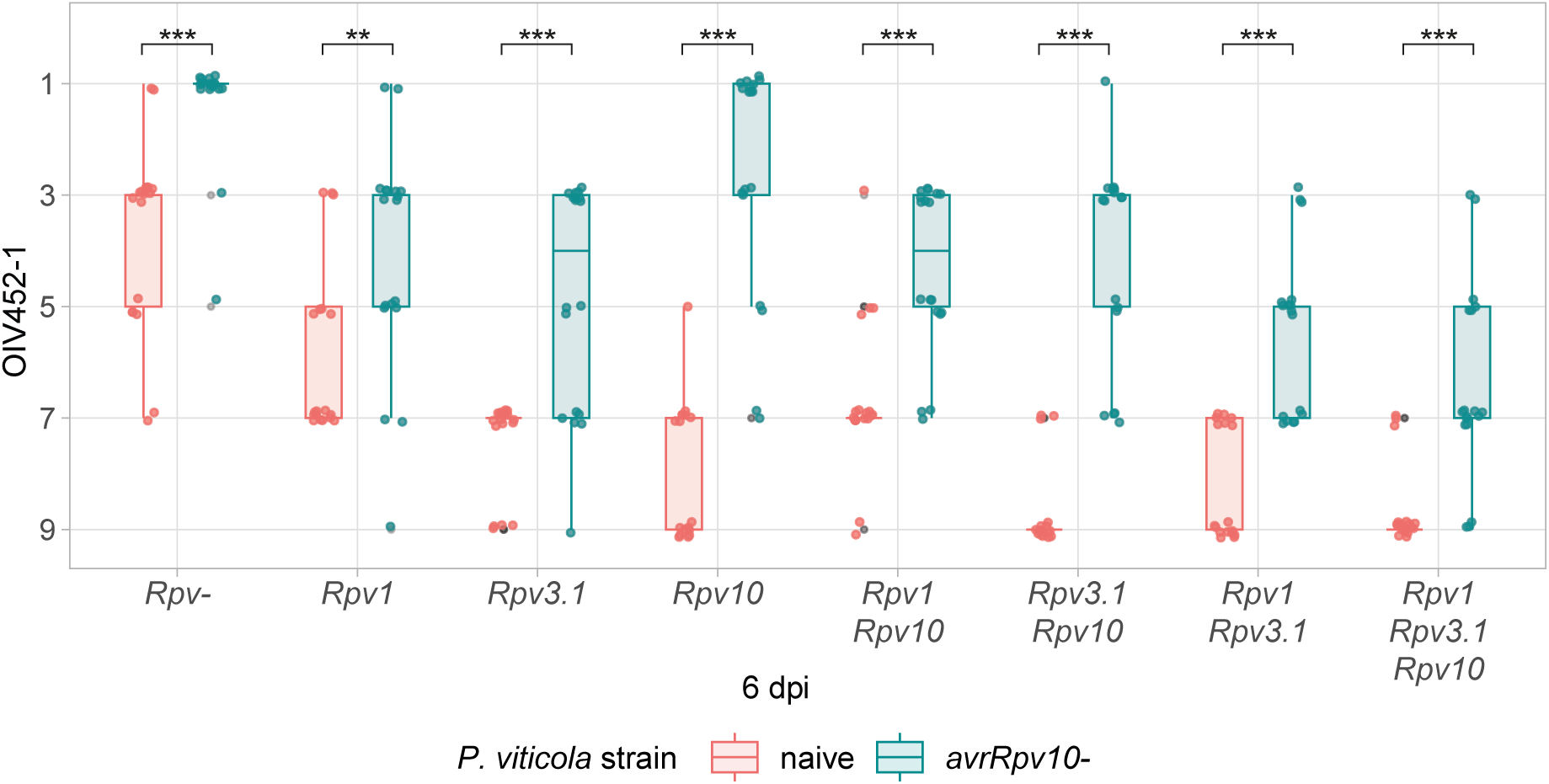
Visual assessment of the infection in the *Rpv* combination-*P. viticola* strains interactions. OIV452-1 scores at 6 dpi for the different *Rpv* combinations challenged with the two *P. viticola* strains (1 = sporulation densely covering the whole disc; 9 = absence of sporulation). Data for two experiments, 3 genotypes/*Rpv* combination and 3 discs /genotype/strain are displayed. Comparisons refer to t-test between *P. viticola* strains infection on the same *Rpv* combination (p value < 0.05 = *, < 0.01 = ** and < 0.001 = ***). Detailed data for the statistical analysis are available in Supplementary File S4b.

**Table 1.** Visual outcomes of the *Rpv* combination-*P. viticola* strain interactions. Mean values for the OIV452-1, *P. viticola* sporulation incidence, *P. viticola* sporulation latency period and plant necroses incidence at 6 dpi for the 16 *Rpv* combination - *P. viticola* strain interactions studied (3 genotypes/*Rpv* combination and 3 leaf discs/genotype/strain for two experiments). Detailed data for the experiments are available in Supplementary File S4a.

| *Rpv* combination | *P. viticola* strain | OIV452-1 | Sporulation incidence | Latency period | Necrosis incidence |
| --- | --- | --- | --- | --- | --- |
| *Rpv*- | naive | 3.67 | 100.00% | 4.00 | 38.89% |
| *Rpv*- | *avrRpv10*- | 1.33 | 100.00% | 3.89 | 5.56% |
| *Rpv1* | naive | 5.89 | 100.00% | 4.11 | 94.44% |
| *Rpv1* | *avrRpv10*- | 4.11 | 94.44% | 3.83 | 61.11% |
| *Rpv3.1* | naive | 7.44 | 77.78% | 4.07 | 88.89% |
| *Rpv3.1* | *avrRpv10*- | 4.78 | 94.44% | 4.12 | 61.11% |
| *Rpv10* | naive | 7.94 | 47.06% | 4.88 | 94.12% |
| *Rpv10* | *avrRpv10*- | 2.56 | 100.00% | 3.94 | 16.67% |
| *Rpv1Rpv10* | naive | 6.67 | 88.89% | 4.88 | 100.00% |
| *Rpv1Rpv10* | *avrRpv10*- | 4.33 | 100.00% | 3.50 | 88.89% |
| *Rpv3.1Rpv10* | naive | 8.65 | 17.65% | 6.00 | 82.35% |
| *Rpv3.1Rpv10* | *avrRpv10*- | 4.18 | 100.00% | 3.82 | 100.00% |
| *Rpv1Rpv3.1* | naive | 8.18 | 41.18% | 5.14 | 82.35% |
| *Rpv1Rpv3.1* | *avrRpv10*- | 5.47 | 100.00% | 4.12 | 88.24% |
| *Rpv1Rpv3.1Rpv10* | naive | 8.67 | 16.67% | 5.75 | 72.22% |
| *Rpv1Rpv3.1Rpv10* | *avrRpv10*- | 6.44 | 83.33% | 4.06 | 94.44% |

The infection with the *P. viticola* naive strain yielded visible necrosis from 3 dpi, with a high frequency on the interactions involving at least one effective *Rpv* (on average the 88% of discs), and a low frequency on the susceptible combination *Rpv-* (34% of discs). Pathogen sporulation started at 4 dpi and increased from 4 to 6 dpi, varying in severity and incidence according to the *Rpv* combination (Fig. 1; Table 1). The OIV452-1 at 6 dpi described susceptibility to *P. viticola* for *Rpv-*, which was always characterized by a severe pathogen sporulation (scores around 3), and partial resistance for single *Rpv1*, *Rpv3.1* and *Rpv10* with lower severity of sporulation (scores between 6 and 8; Fig. 2; Table 1). Accordingly, ANOVA and pairwise comparisons (PWC) between *Rpv*, identified a significant effect of *Rpv1*, *Rpv3.1* and *Rpv10* on the overall offspring resistance and showed an averaged resistance level *Rpv10≥Rpv3.1>Rpv1* (Table 2; Supplementary File S4c-4d). The *Rpv3.1Rpv10, Rpv1Rpv3.1* and *Rpv1Rpv3.1Rpv10* pyramiding combinations enhanced and stabilised the resistance with respect *Rpv1* and *Rpv3.1* single loci, resulting in less frequent, delayed (5-6 dpi) and reduced pathogen sporulation (scores typically around 7-9; Fig. 1; Table. 1; Supplementary File S4d). With respect to *Rpv10*, pyramiding gains were more limited and not significant, in particular for *Rpv1Rpv10* combination, which showed intermediate level of infection between *Rpv10* and *Rpv1* (Table. 1; Supplementary File S4d).

**Table 2.** ANOVA results for visual assessment of *Rpv* combination*-P. viticola* strains interactions. Results from strain-independent ANOVA conducted with the OIV452-1 scores at 6 dpi, and including *Rpv1*, *Rpv3.1* and *Rpv10* presence (two levels with complete cross interactions), and the Experiment, as factors. Data for 3 genotypes/*Rpv* combination and 3 discs /genotype/strain of two experiments were considered. Detailed data for the statistical analysis are available in Supplementary File S4c.

| ANOVA | Factors | df | F value | p. value |
| --- | --- | --- | --- | --- |
| OIV452-1 scores at 6 dpi for the *P. viticola* strain naive | *Rpv1* | 1 | 4.338 | 0.039 |
|  | *Rpv3.1* | 1 | 117.730 | <0.001 |
|  | *Rpv10* | 1 | 69.684 | <0.001 |
|  | Exp | 1 | 4.893 | 0.029 |
| OIV452-1 scores at 6 dpi for the *P. viticola* strain *avrRpv10*- | *Rpv1* | 1 | 41.723 | <0.001 |
|  | *Rpv3.1* | 1 | 54.647 | <0.001 |
|  | *Rpv10* | 1 | 2.633 | 0.107 |
|  | Exp | 1 | 11.253 | 0.001 |

*Rpv-* and *Rpv10* genotypes when challenged with the virulent strain *avrRpv10-,* exhibited both susceptible phenotypes with high sporulation rates and severity (OIV452-1 scores of 1 and 3) and reduced incidence of necrosis (about 11% of the discs; Fig. 1; Table 1). Additionally, the virulent strain displayed a general higher aggressiveness compared to the naive strain, displaying a fast, more frequent and pronounced sporulation on *Rpv-*, single *Rpv1* and *Rpv3.1*, and pyramided *Rpv* (scores between 4 and 6 for all resistance interactions; Fig. 1-2; Table 1). ANOVA of OIV452-1 scores at 6 dpi identified as significant the effect of both *Rpv1* and *Rpv3.1* on the offspring resistance, while *Rpv10* presence was not identified as a significant factor in the analysis (Table 2; Supplementary File S4c). PWC between *Rpv* combinations confirmed similar severity of the infection at 6 dpi on *Rpv1* and *Rpv3.1* single loci, and, due to *Rpv10* breakdown, the severity of the infection on *Rpv1Rpv10* and *Rpv3.1Rpv10* combinations did not differ from single loci (Supplementary File 4d). Only the pyramiding of *Rpv1Rpv3.1* and *Rpv1Rpv3.1Rpv10* significantly increased the resistance level with respect other single and pyramided *Rpv* combinations (details of comparisons in Supplementary File 4d). This last result also suggested a residual effectiveness of *Rpv10* in controlling the *P. viticola avrRpv10-* strain.

The ANOVA analysis revealed a significant effect of the experiment on both strains (Table 1). According to data distribution a lower rate of infection occurred in the second experiment (Supplementary File 4b). However, the infection progression and relative level of infection between *Rpv* combinations and *P. viticola* strains were consistent in the two replicates (Supplementary File 3; 4a). Thus, the experiments were both validated and considered equally representative of the studied conditions, and the first experiment was chosen to perform in-depth metabolomics analysis.

### Assessment of *P. viticola* metabolic biomarkers

The metabolomic analysis focused on 12 *P. viticola*-specific biomarkers characterized previously [18] (Supplementary File S5a), including: eicosapentaenoic acid (EPA) and three of its derivatives (eicosapentaenoyl-glycerol – EPG, dieicosapentaenoyl-glycerol – DEPG and trieicosapentaenoylglycerol – TEPG), arachidonic acid (AA) and three of its derivates (arachidonoyl-glycerol – AG, diarachidonoyl-glycerol – DAG and triarachidonoyl-glycerol – TAG), and four ceramides (Cer(d16:1/16:0), Cer(d16:1/18:0), Cer(d16:1/20:0) and Cer(d16:1/22:0)). The compounds were quantified by an optimized analysis protocol based on liquid chromatography-based mass spectrometry (LC-MS) and targeted metabolomics (details in Material and methods section). All the 16 *Rpv* loci-*P. viticola* strain interactions were studied at 6-12 hours post inoculation (hpi) and at 1-2-3 dpi to investigate the early phase of infection, and at 6 dpi to precisely quantify the infection (3 infected discs/genotype/strain/time). All the *P. viticola* biomarkers were found in visually high-infected discs at 6 dpi, but only EPA, AA, AG, Cer(d16:1/18:0), Cer(d16:1/20:0) and Cer(d16:1/22:0) were consistently identified in the different studied interactions and in the early phase of the infection (at least in the 70% of the samples; Supplementary File S5b). Finally, all the better-detected biomarkers except AG resulted highly correlated with each other (R^2^ = 0.87 - 0.98) and with the OIV452-1 scores (R^2^ = 0.82 to 0.91; Supplementary File S5c-5d) and were retained as informative features for the quantitative evaluation of the *Rpv*-*P. viticola* interactions (Table 3). In particular, Cer(d16:1/22:0) showed the earliest and largest signal variation in the different studied conditions (Fig. 3a; Supplementary File S6)

**Figure 3.**
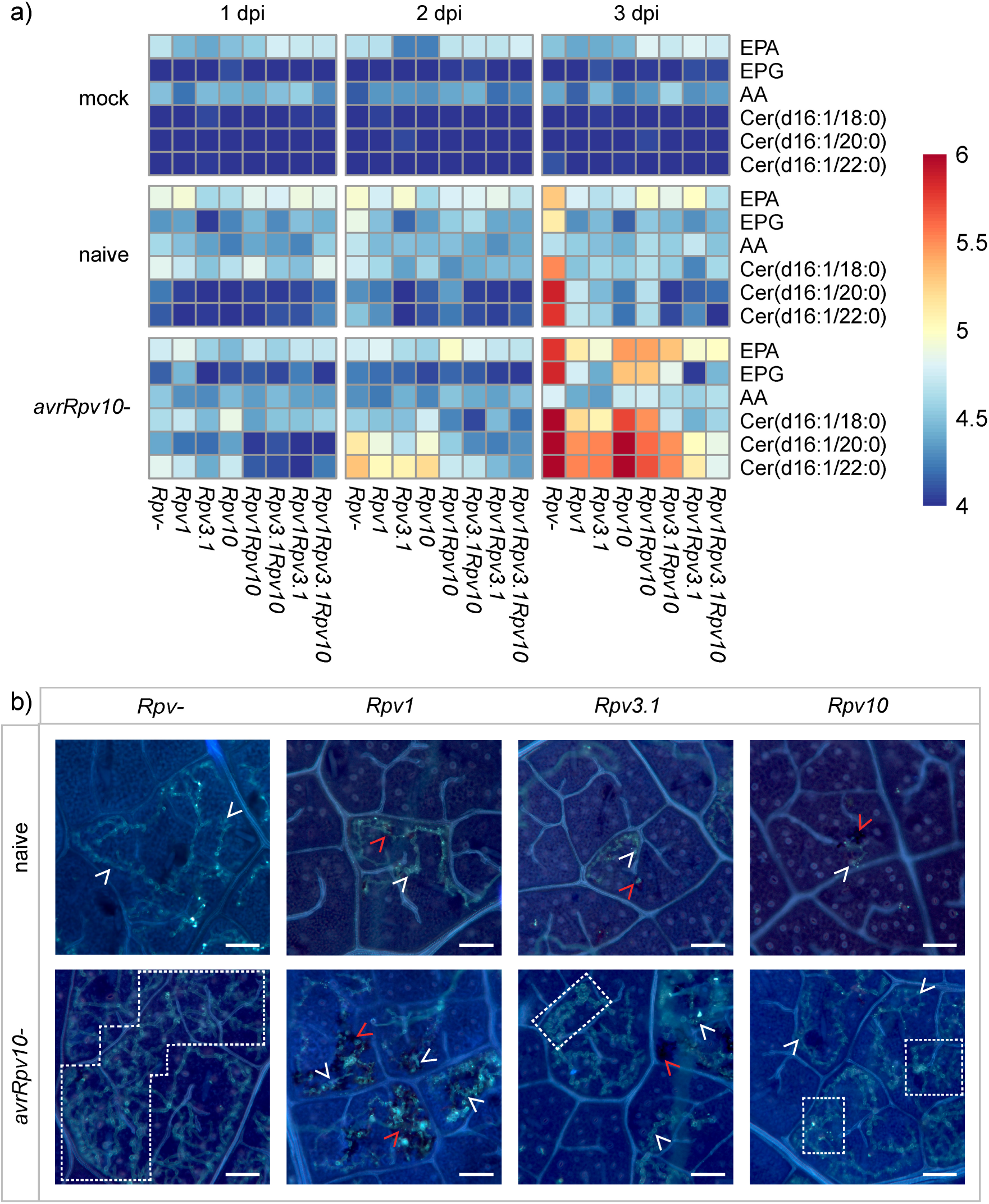
Assessment of the early phase of the infection in the *Rpv* combination-P. *viticola* strain interactions. a) The heatmap shows the mean peak area (arbitrary unit -AU) for the six most informative *P vitico/a-specific* metabolic biomarkers (right labels) recorded in the discs of the different *Rpv* combinations mock-inoculated or infected with the two *P viticola* strains (naive and *avrRpv10-)* at 1-2-3 dpi. Data for one experiment, 3 genotypes/Rpv combination and 3 discs/genotype/strain/dpi are displayed. b) Representative microscopic images after the staining of *P vitico/a* mycelium by aniline-blue for the non-resistant (Rpv-) and single *Rpv* combinations inoculated with the two *P vitico­*/a strains at 3 dpi: the white arrows indicate *P vitico/a* mycelium, the white dashed-square the area with dense and overlapping layers of mycelium, and red arrows sites where *P vitico/a* infection is associated to plant necrotic response (scale bar 100 µm).

**Table 3.** *P. viticola* metabolic biomarkers followed by targeted analysis. Information on the *P. viticola* metabolic biomarkers consistently identified throughout the experiment and informative for the quantification of *P. viticola* infection/biomass: compound name and acronym, ion formula, mass (m/z) and retention time (RT) utilized for the quantification. The complete list of *P. viticola* biomarkers investigated is available in Supplementary File S5a.

| Compound | Acronym | Ion formula | m/z | RT(s) | Identifier |
| --- | --- | --- | --- | --- | --- |
| Eicosapentaenoic acid | EPA | C20H30O2 | 303.23169 | 118.20 | M303T118 |
| Eicosapentaenoyl-glycerol | EPG | C23H37O4 | 377.26855 | 106.20 | M377T106 |
| Arachidonic acid | AA | C20H32O2 | 305.24741 | 126.60 | M305T127 |
| Cer(d16:1/18:0) | Cer(d16:1/18:0) | C34H66NO2 | 520.50842 | 163.20 | M521T163 |
| Cer(d16:1/20:0) | Cer(d16:1/20:0) | C36H70NO2 | 548.53986 | 168.00 | M549T168 |
| Cer(d16:1/22:0) | Cer(d16:1/22:0) | C38H74NO2 | 576.57129 | 171.60 | M577T172 |

#### Rpv loci-P. viticola strain interactions in the early phase of the infection

Metabolomics did not show significant variations in the accumulation of *P. viticola*-specific compounds before 1 dpi (Supplementary File S6). According to pathogen biomarkers abundances fold-changes (fc), the *P. viticola* infection increased between 2-50 folds from 1 to 3 dpi, depending on the *Rpv*-*P. viticola* strain interaction considered. The *P. viticola avrRpv10-* strain showed higher biomarkers accumulations from 2 dpi on, comparisons at 3 dpi clearly demonstrating an enhanced aggressiveness in the early phase of the infection compared to the naive strain on all *Rpv* combinations (e.g., 4-64 fc for the quantified ceramides; Fig. 3-4a; Supplementary File S7a-8a).

**Figure 4.**
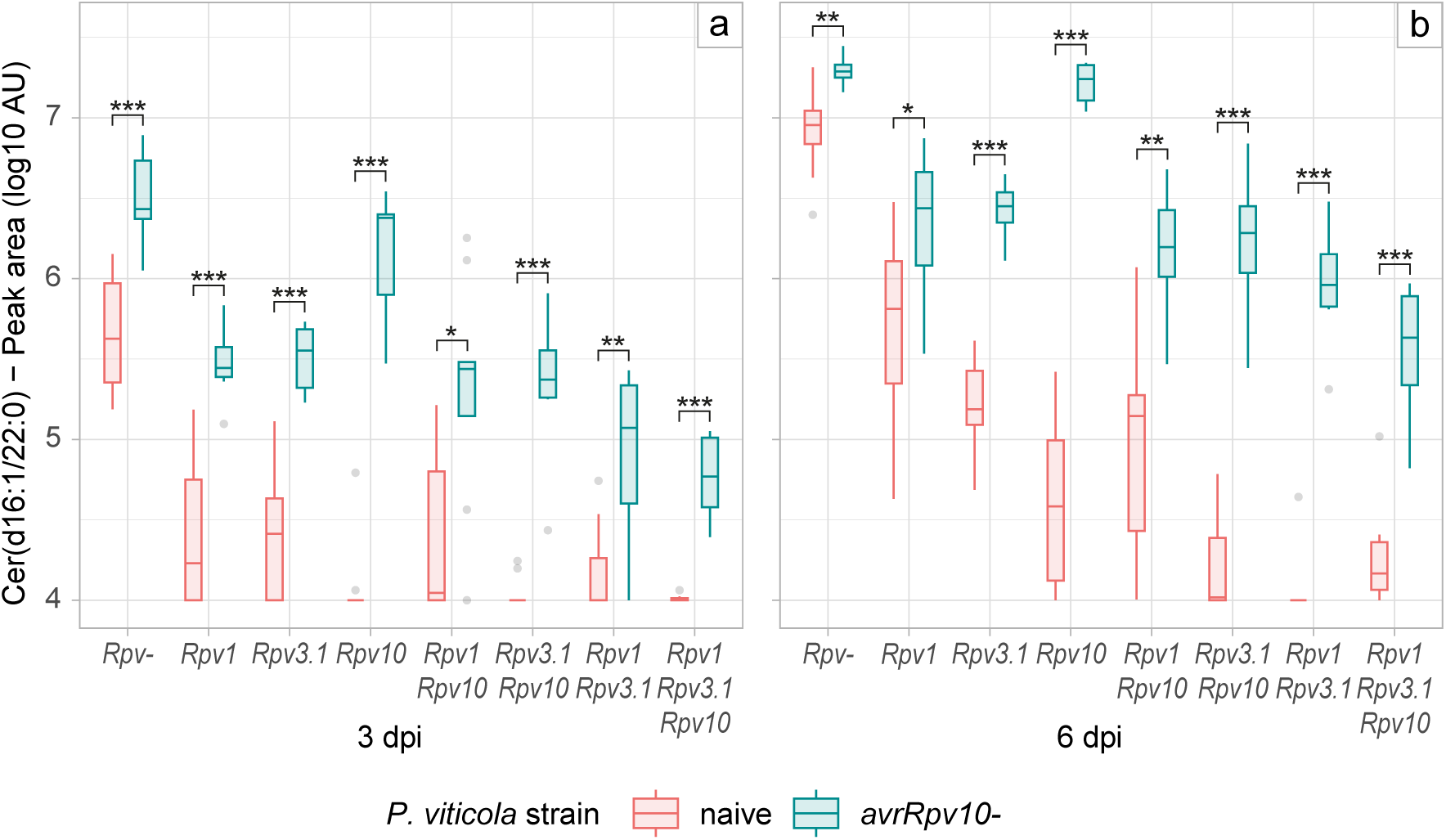
Metabolomics-based quantification of infection in the in the *Rpv* combination-*P. viticola* strain interactions. *P. viticola* metabolic biomarker Cer(d16:1/22:0) abundances at 3 (a) and 6 (b) dpi for the different *Rpv* combinations challenged with the two *P. viticola* strains. Abundances are displayed as log10 of the peak area arbitrary unit (AU). Data for one experiment, 3 genotypes/*Rpv* combination and 3 discs/genotype/strain/dpi are displayed. Comparisons refer to t-test between *P. viticola* strains infection on the same *Rpv* combination (p value < 0.05 = *, < 0.01 = ** and < 0.001 = ***). Detailed data for the statistical analysis are available in Supplementary File S8a.

Microscopy observations of *P. viticola* aniline blue-stained mycelium supported the results inferred from the analysis metabolic biomarkers, confirming significant differences in the pathogen colonisation rate at 3 dpi on the three single Rpv loci studied. For the *P. viticola* naive strain, an important colonisation with ramified pathogen mycelium (white arrows in Fig. 3b) was observed on *Rpv-*, limited hyphae growth was associated to plant necrosis (red arrows in Fig. 3b) on *Rpv1* and *Rpv3.1*, and stopped infections occurred on *Rpv10* (Fig. 3b). For the *P. viticola avrRpv10-* strain, dense and overlapping layers of mycelium was observed on both *Rpv-* and *Rpv10*, and important colonisations with ramified mycelium occurred on *Rpv1* and *Rpv3.1* despite a wide presence of plant necrosis (Fig. 3b).

Limitation of *P. viticola* infection at 3 dpi (e.g., 8-32 fc for ceramides accumulation compared to *Rpv-*) were associated with *Rpv-*mediated resistance (Fig. 3-4a; Supplementary File S7b). For the naive strain, the ANOVA of Cer(d16:1/22:0) data identified the presence of *Rpv3.1* and *Rpv10* as significant on the offspring resistance, while *Rpv1* contribution was not identified (Table 4; Supplementary File S8b), probably due to the limited differences between *Rpv* combinations, as described by the data distribution and the PWC (Fig. 4a; Supplementary File S8c). For the *avrRpv10-* strain, the globally higher rate of infection allowed to better differentiate the resistance mediated by the different *Rpv* combinations (Fig. 4a). The ANOVA of Cer(d16:1/22:0) data identified both the *Rpv1* and *Rpv3.1* effects on the offspring resistance (Table 4; Supplementary File S8b), and PWC evidenced a lower rate of infection also between *Rpv1Rpv3.1* and *Rpv1Rpv3.1Rpv10* compared to the other resistance combinations (Fig. 4a; Supplementary File S8c). Intriguingly, *Rpv10* as a single locus induced a reduced pathogen biomass at 2-3 dpi compared to *Rpv-*. Similarly, *Rpv1Rpv10* or *Rpv3.1Rpv10* combinations exhibited a reduced pathogen biomass compared to *Rpv1* or *Rpv3.1* single loci, although none of these differences were statistically significant (Supplementary File 8b-8c).

**Table 4.** ANOVA results for the metabolomics-based quantification of infection in the *Rpv* combination*-P. viticola* strain interactions. Results from strain-independent ANOVA conducted with *P. viticola* biomarker Cer(d16:1/22:0) abundances (log10 of the peak area arbitrary unit AU) at 3 and 6 dpi, and including *Rpv1*, *Rpv3.1* and *Rpv10* presence (two levels with complete cross interactions) as factors. Data for one experiment, 3 genotypes/*Rpv* combination and 3 discs/genotype/strain/dpi were considered. Detailed data for the statistical analysis are available in Supplementary File S8b.

| **ANOVA** | **Factors** | **df** | **f value** | **p value** |
| --- | --- | --- | --- | --- |
| Cer(d16:1/22:0) at<br>3 dpi for the *P.*<br>*viticola* strain naive | *Rpv1* | 1 | 2.218 | 0.141 |
|  | *Rpv3.1* | 1 | 11.617 | 0.001 |
|  | *Rpv10* | 1 | 29.890 | <0.001 |
| Cer(d16:1/22:0) at<br>3 dpi for the *P.*<br>*viticola* strain<br>*avrRpv10*- | *Rpv1* | 1 | 49.592 | <0.001 |
|  | *Rpv3.1* | 1 | 38.575 | <0.001 |
|  | *Rpv10* | 1 | 3.350 | 0.072 |
| Cer(d16:1/22:0) at<br>6 dpi for the *P.*<br>*viticola* strain naive | *Rpv1* | 1 | 5.674 | 0.020 |
|  | *Rpv3.1* | 1 | 47.746 | <0.001 |
|  | *Rpv10* | 1 | 36.341 | <0.001 |
| Cer(d16:1/22:0) at<br>6 dpi for the *P.*<br>*viticola* strain<br>*avrRpv10*- | *Rpv1* | 1 | 96.516 | <0.001 |
|  | *Rpv3.1* | 1 | 79.240 | <0.001 |
|  | *Rpv10* | 1 | 7.063 | 0.010 |

#### Rpv loci-P. viticola strain interactions in the later phase of infection

*P. viticola* infection for the naive and *avrRpv10-* strains increased consistently from 3 to 6 dpi on all the studied *Rpv* combinations, except for the naive strain on the most effective pyramided combinations *Rpv3.1Rpv10*, *Rpv1Rpv3.1* and *Rpv1Rpv3.1Rpv10* (Fig. 4). The amplitude of some differences in *P. viticola* biomarker abundances between *Rpv* combinations or *P. viticola* strains at 3 and 6 dpi changed. For example, the differences between *P. viticola* strains on *Rpv-* and *Rpv1,* and between *Rpv-* and *Rpv10* for the *avrRpv10-* strain, were more important at 3 dpi than at 6 dpi (Fig. 4), while the difference between *Rpv1* and *Rpv3.1* for the naive strain were higher at 6 dpi than at 3 dpi. This suggests that some specificities of *Rpv*-*P. viticola* interactions could be better visible in the early or in the late phase of infection. Although the majority of outcomes at 3 and 6 dpi remained strongly in agreement and defined a general relationship between the early pathogen colonization and the later sporulation (Fig. 4; Supplementary File S6).

Biomarker data at 6 dpi confirmed OIV452-1 observations, but they provided a more objective and quantitative perspective on *Rpv*-mediated resistance and *P. viticola* aggressiveness. Compared to *Rpv-*, *P. viticola* ceramides showed a significant infection reduction of 6-9 fc on *Rpv1* for both strains, of 47 and 6 fc on *Rpv3.1* for the naive and *avrRpv10-* strains, respectively, and of 100 fc on *Rpv10* for the naive strain (Fig. 4b; Supplementary File S7b-S8c). On the other hand, for the *P. viticola avrRpv10-* strain, the biomarkers highlighted a greater infection of 2-3 fc*-* on *Rpv-* and *Rpv1*, and of 16 fc on *Rpv3.1*, suggesting a partial virulence or an adaptation of the *avrRpv10-* strain towards *Rpv3.1* (Fig. 4b; Supplementary File S7a-S8a). The *Rpv*-pyramiding effects were also precisely quantified, as evidenced for *Rpv1Rpv3.1* combination, which, in comparison with single loci, reduced the pathogen biomass of 10-50 fc for the naive strain, and of 2-2.5 fc for the *avrRpv10-* strain (Fig. 4b; Supplementary File S7b-S8c). Interestingly, the ANOVA of Cer(d16:1/22:0) at 6 dpi identified a significant resistance effect for *Rpv10* against the *avrRpv10-* strain (Fig. 5a; Table 4; Supplementary File S8b), suggesting a general residual resistance effect of the locus. In particular, the *Rpv10* residual effect significantly reduced the infection of 2 folds when pyramided in the *Rpv1Rpv3.1Rpv10* combination (Fig. 5b; Supplementary File S8c).

**Figure 5.**
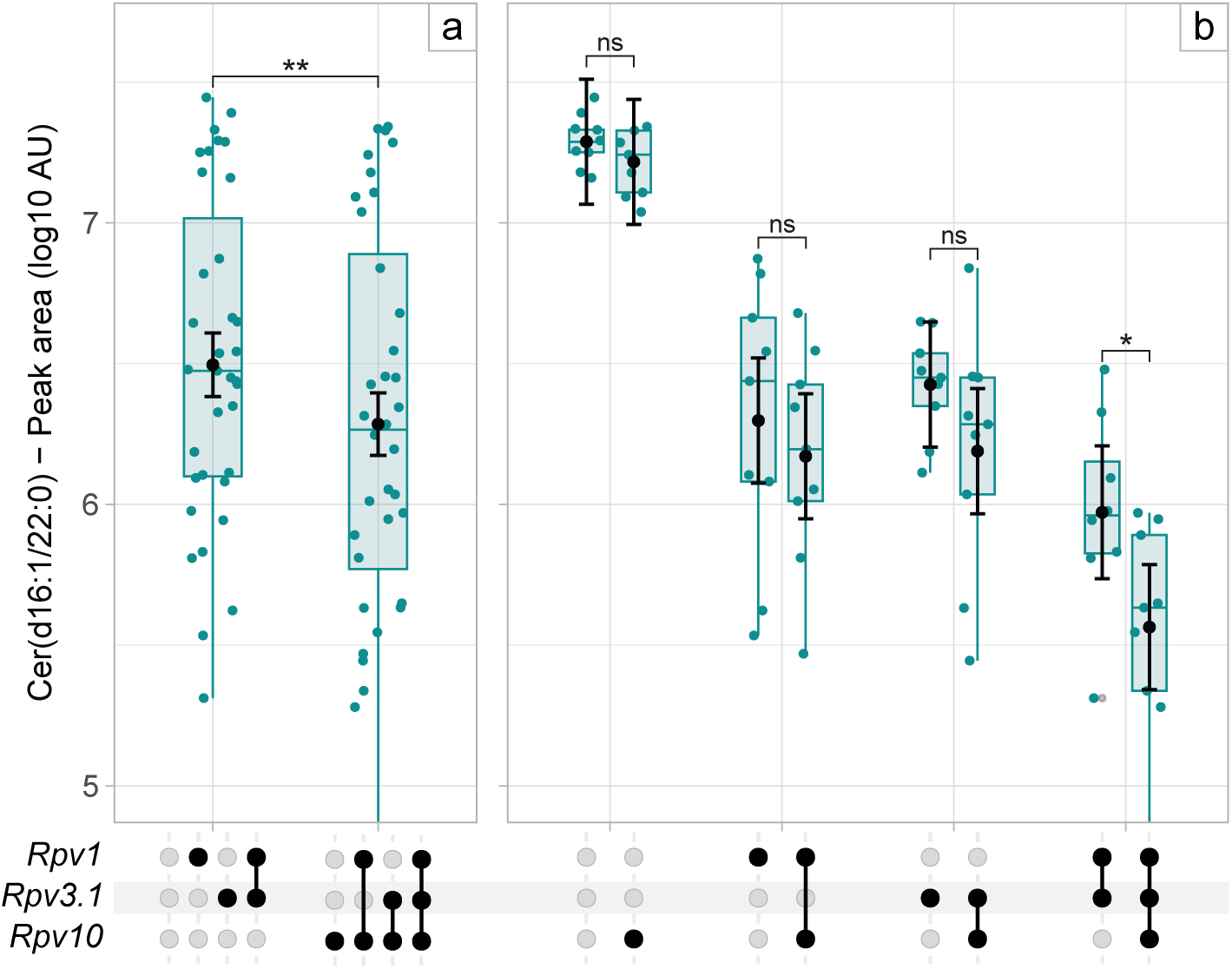
*Rpv10* effect against the virulent *P. viticola* strain *avrRpv10-*. ANOVA (a) and PWC (b) for *P. viticola* biomarker Cer(d16:1/22:0) abundances detected at 6 dpi in the different *Rpv* combinations infected with the *P. viticola* strain *avrRpv10-* (*Rpv* combinations considered in the comparisons are indicated by black dots-lines on the x-axis). Abundances are displayed as log10 of the peak area arbitrary unit (AU) in the box plots, while the black dots and error bars indicate the estimated marginal means and standard errors for each comparison group (Tukey’s tests p value < 0.05 = *, < 0.01 = ** and < 0.001 = ***). Data for one experiment, 3 genotypes/*Rpv* combination and 3 discs/genotype/strain/dpi are displayed. Detailed data for the statistical analysis are available in Supplementary File S8b-8c.

## Discussion

Several *Rpv* loci and virulent *P. viticola* strains have been described [7,12,13]. However, former studies usually relied on a limited number of unrelated grape varieties and visual descriptors for the characterization of *Rpv*-mediated resistance and *P. viticola* aggressiveness [11,12,21], with potential influences of varietal genetic background [21,22] and phenotyping technical limitations. In our study, we address these issues by examining all the possible combinations of single and pyramided *Rpv* in a full-sib offspring, using *P. viticola* metabolic biomarkers to provide an objective and accurate quantification of the infection. This approach highlights the early and late effects of *Rpv-*mediated resistance, as well as the impact *P. viticola* aggressiveness on infection dynamics, revealing positive interactions among pyramided loci, even for the a defeated *Rpv,* and different level of adaptations of the virulent *P. viticola* strains towards the host genetic resistance.

### Single *Rpv* showed specific interactions when inoculated with different *P. viticola* strains

*Rpv* loci have been associated with different resistance levels, ranging from total to strong, intermediate, or weak partial resistances [7]. In our study, the resistance conferred by *Rpv1*, *Rpv3.1* and *Rpv10* was partial and changed in response to the two *P. viticola* strains tested. We observed that the efficacy of *Rpv* followed the ranking *Rpv10*>*Rpv3.1*>*Rpv1* with the naive *P. viticola* strain. A different efficacy was observed with the *avrRpv10-* strain, for which Rpv*1*=*Rpv3.1* and *Rpv10=Rpv-*, suggesting specific interaction of the strains with the studied *Rpv*.

On *Rpv*- we observed for the *P. viticola avrRpv10-* strain, compared to the naive strain, a greater infection and mycelium growth at 3 dpi, and a greater sporulation at 6 dpi, which suggested an enhanced basal aggressiveness of the *avrRpv10-* strain. The existence of variations in pathogen aggressiveness is a key factor in pathogen adaptation [11,23], and higher *P. viticola* basal aggressiveness has been described in isolates adapted to cope with resistant varieties [11,23]. Thus, an enhanced aggressiveness can be considered as a typical characteristic of *P. viticola* strains with specific adaptations towards *Rpv* loci, and it could be essential for deploying certain virulence mechanisms.

*Rpv10* provided a strong partial resistance towards the *P. viticola* naive strain. However, large resistance variability was associated to *Rpv10* when challenged with a single *P. viticola* strain [24], revealing possible impacts of the varietal genetic background [25] and the environment [24,26] on the *Rpv10* effectiveness. In our study, the stability of resistance phenotypes probably resulted from the structure of the grapevine population and the controlled laboratory conditions. The interaction between *Rpv10* and the *avrRpv10-* strain resulted in susceptible phenotypes, characterised by high sporulation and low frequency of necrosis, in agreement with previous studies on *Rpv* breakdown [12,27]. The described *P. viticola avrRpv10-* strain represents an additional pathotype among those identified in Europe, as the strains virulent towards *Rpv10* were usually described as virulent towards *Rpv3.1* as well [12,21,27]. However, *Rpv10* breakdown has been associated to a single genomic admixture event localized in Central Europe [27]. Thus, the virulence of *avrRpv10-* strain should be link to a dominant locus and to the suppressor activity of specific pathogen effectors [27]. On the other hand, the *P. viticola avrRpv10-* strain showed only a partial adaption on *Rpv3.1* compared to other strains overcoming *Rpv10,* posing questions on the original pathotype of the newly identified *P. viticola* population and the specific adaptation of the *avrRpv10-* strain.

Different level of partial resistance were described for *Rpv3.1*, and were associated to weak, intermediate or strong sporulation of *P. viticola* [7,12,21]. In our study, *Rpv3.1* was effective in controlling the *P. viticola* naive strain, while the *avrRpv10-* strain showed an enhanced sporulation on *Rpv3.1*-carrying combinations. Different levels of *P. viticola* adaptation to *Rpv3.1* were associated with specific changes in the assortment of a set of pathogen effectors [28]. The *P. viticola* virulence on *Rpv3.1* is a recessive character determined by the loss of a couple of effectors [28]. The variability in the *Rpv3.1-*mediated resistance recorded in our study can be probably depend from different patterns characterizing the effector set of the two *P. viticola* strains used here.

Our findings indicated a similar effectiveness of *Rpv1* in controlling the *P. viticola* naive strain and the *avrRpv10-* strain, in agreement with other screenings of *P. viticola* strains [12,21]. Pathogens usually quickly adapt to qualitative plant resistances, which exert strong selection pressures on pathogen populations [29]. The collected evidences indicate that in grapevine the widely used and strong *Rpv* loci, such as *Rpv3.1*, have been rapidly defeated [12,21,28]. On the other hand, the weaker and less utilized locus *Rpv1* has retained its effectiveness for longer [12,21], although climatic conditions highly favourable to *P. viticola* have recently determined localized *Rpv1* breakdown [30].

### Toward reasoned pyramiding to improve grapevine resistance

Plant genetic resistances are limited resources and their breakdown is an irreversible gain for pathogens, that compromise resistance breeding in the long-term [24,31]. Therefore, *Rpvs* should be used in pyramiding in viticulture, as this strategy is the most appropriate for the deployment of grapevine genetic diversity [24,31], in order to increase the resistance efficacy, stability and durability [14,32,33].

Our study confirmed the efficacy of pyramiding, and showed that the stacking of two functional *Rpvs* was already highly effective against the naive *P. viticola* strain, severely delaying and reducing the pathogen infection. Pyramiding was less effective when *Rpv1*, *Rpv3.1* and *Rpv10* were challenged with the *avrRpv10-* strain, adapted to cope with resistant varieties thanks to a higher basal aggressiveness and *Rpv*-specific adaptations. However, we still observed the highest resistance levels in the *Rpv* combination carrying three loci, identifying a significant residual benefit from the defeated *Rpv10*. This results is in agreement with the observations made for the pyramiding of the defeated *Rpv10* and *Rpv12* in another grapevine population [24], and the outcomes linked to defeated resistance genes in other plant-fungal pathosystems [34,35]. The cumulation of defeated loci has been demonstrated as a strategy to gain moderate-to-high levels of resistance [36], but in grapevine the stacking of defeated *Rpvs* has instead yielded negative effects on the final level of resistance in some instances [24]. Nevertheless, residual quantitative resistance effects should be monitored over time and across different sites [35]. Indeed, *P. viticola* showed both common and distinct genetic mutations in its convergent adaptation to *Rpv,* which differentially affected the pathogen virulence [37].

The number of resistance genes is not the only factor affecting pyramiding outcomes [38]. Among the multiple pyramiding combinations tested, the *Rpv1Rpv10* combination showed neutral (towards *Rpv1*) or negative (towards *Rpv10*) effects on resistance with the *P. viticola* naive strain. Undesired effects in loci pyramiding have been described in other in crops [38,39], and were identified in grapevine for the combination *Rpv3.1Rpv10* [21] in some varieties, and for *Rpv1Rpv10* when challenged with an *avrRpv10-* strain [24]. However, in our study the *Rpv3.1Rpv10* combination was beneficial in enhancing the resistance towards the naive strain, and *Rpv10* did not compromise *Rpv1* resistance towards the *avrRpv10-* strain. Despite the unpredictability of negative *Rpv* interactions, in-depth studies are needed to confirm and explain the causes of these interactions, as they pose a threat to the resistance efficacy and durability [24].

Finally, resistance breeding should include both the pyramiding of compatible NLR genes [24] and alternative sources of quantitative resistance [14,32,33], such as constitutive (e.g., structural barriers and secondary metabolites) and other induced defences (e.g., pathogenesis-related proteins and cell wall thickening) [32,33]. Quantitative resistance is still underutilized in grapevine breeding, but it could be highly beneficial, as it could have additive resistance effects, be effective against multiple strains, and provide protection against the breakdown of major genes [33]. However, the quantitative nature of such resistance, which is often based on multiple genes with limited effects, makes it difficult to study and introduce in selection programs [32,33].

### From ephemeral to effective plant resistance to pathogens

The early stages of *P. viticola* infection have typically been described using microscopy and the attempts to quantify the early resistance effects have resulted in laborious and low resolution protocols [13,40,41], as the detection and classification of *P. viticola* spores growth stages [40,41]. Recent advancements in the early assessment of *P. viticola* infection have involved the application of real-time PCR [42,43] and metabolomics [18]. In our study, considerable variations in *P. viticola* biomarkers abundances were observed between *Rpv* combinations and strains from 1 dpi. The observations indicated that the effects of resistance loci and pathogen aggressiveness manifested at the onset of the infection. However, some resistance effects were ephemeral or became less evident in later stages of infection, as the limited effect of single *Rpv10* towards the *avrRpv10-* strain. This information can be of great help to understand the mechanisms of plant-pathogen interactions. For example, the detection of early effects of *Rpv10* on the *P. viticola avrRpv10-* strain may corroborate the hypothesis that, as a single locus, *Rpv10* activates early resistance mechanisms, but these are subsequently suppressed by the pathogen [27], which is then able to recover from the initial delay with exponential growth. In contrast, the defeated *Rpv10* has proved to be more effective in pyramiding, but the data collected did not permit to determine whether *Rpv10* interacts directly with the other *Rpv*, for an early and/or stronger resistance response, or if the initial *Rpv10-*dependent delay in pathogen progression allows a more effective deployment of resistance mechanisms mediated by other pyramided *Rpvs*. This finding also suggests that ephemeral resistances during the early stages of infection can provide valuables effects when combined with other resistance factors. Consequently, we propose for future research on plant resistance to encompass evaluations in both the early and late phases of infection, as early interactions in pyramiding may provide significant benefits for efficient pathogen control. This aspect is of particular interest in characterization of alternative sources of quantitative resistance, which often depends on several loci with effects of variable amplitude [32,33].

### Metabolic biomarkers as an effective strategy to characterize plant-pathogen interactions

The assessment of *P. viticola* infection on leaf discs has traditionally been conducted through visual variables, taking into account the severity of the disease, the density of the pathogen structures, and the count of sporangia as the primary criteria [7,12]. In our study, we employed LC-MS to quantify *P. viticola* metabolic biomarkers and subsequently inferred the rate of infection. In comparison to visual assessment, biomarkers were objective, had a continuous quantification and enhanced precision, sensitivity and dynamics, which permitted to detect variations in infection rate at any level of the visual scoring, including samples that did not display visual symptoms or reached the highest level of the assessment scale. Therefore, metabolomics provided high-resolution results and allowed to better uncover, infer, and communicate the biological considerations on the pathosystem.

Based on the current literature, our utilization of pathogen-specific metabolic biomarkers represents a novel research approach, given that metabolomics approaches have often been focused on the plant metabolome to characterize changes in metabolic pathways linked to the resistance response [44], as evidenced by the studies on grapevine [45,46], and that pathogen biomarkers have been rather used as detection tools [47]. The development of common/standard metabolomics quantification protocols for pathogen-related compounds could represent an opportunity in the quantitative study of plant-pathogen interactions, as some compounds are shared between species or higher taxonomic levels. For instance, EPA, AA [48] and pathogen-specific ceramides [49] have been also associated to *Phytophthora infestans*, the oomycete causing the late blight of potato (*Solanum tuberosum*). Metabolomics also offers the possibility to discriminate pathogen species and strains [50] and investigate fungal metabolisms [51]. Metabolomics is therefore a concrete transversal resource that can be used to both characterize and quantify different aspects of the plants-pathogen interaction, as well as to elucidate the genetic basis and mechanisms of plant resistance or pathogen aggressiveness.

## Materials and Methods

### Plant material

Grapevines carrying *Rpv1*, *Rpv3.1* and *Rpv10* in eight different combinations were selected from a study full-sib population produced and conserved at INRAE - UMR 1131 SVQV (Colmar, France) [52]. Since 2017, the progenies have been grown as grafted plants in 4 l pots in greenhouse, receiving natural light and a nutritive solution [20]. During the vegetative season, shoots were periodically pruned and pests and diseases were managed through the application of sprays every two weeks. For the bioassays, the fifth and sixth leaves from the apex of two actively growing shoots (typically the 1^st^-2^nd^ not shiny leaves) were collected and used to produce 2 cm leaf discs [20].

### Disease evaluation

The plants were phenotyped in laboratory conditions on leaf discs as described in Macia et al., [20]. Two *P. viticola* strains were parallelly spray inoculated (50,000 sporangia/ml) on different plates: the naive, or avirulent, strain was isolated from the susceptible grape cultivar ‘Chardonnay’ [13,20]; and a *Rpv10*-breaking (*avrRpv10-*), or virulent, strain was isolated from the resistant *Rpv10*-carrying cultivar ‘Muscaris’ [24].

Discs were visually evaluated by the OIV-452-1 descriptor [19] as described in Macia et al., [20], from 3 to 6 dpi, using a five-classes scales: 1 = sporulation densely covering the whole disc; 3 = sporulation present in most of the disc area in large patches; 5 = sporulation present in delimited intercostal patches; 7 = sparse sporulation; and 9 = absence of sporulation. For each *Rpv* combination - *P. viticola* strain interaction, the incidence of discs showing necrotic response to the infection was collected, while the data for the OIV452-1 were used to infer *P. viticola* sporulation incidence and the sporulation latency period. A total of 3 genotypes/*Rpv* and 3 discs/strain/genotype were produced and assessed in two experiments.

*P. viticola*-specific biomarkers were extensively quantified through LC-MS at 6 and 12 hpi, at 1, 2 and 3 dpi, and at 6 dpi using the visually assessed leaf discs. A total of 3 discs/ genotype/strain/time, as well as 2 mock (water)-inoculated discs/ genotype/ time were produced, sampled and stored in 2 ml tubes at -20°C until the start of metabolomics analysis. A randomization scheme for 72 discs was replicated for each sampling condition (strain/time). Metabolomics-based assessment at 3 dpi was corroborated by investigating *P. viticola* development at by aniline-blue staining as described in Trouvelot et al. [53]. Observations were carried out using a Zeiss Axio Imager M2 (Zeiss, Oberkochen, Germany) in epifluorescence microscopy under UV (λexc 340 nm, λem380 nm, stop filter LP 430 nm).

Two consecutive experiments were carried out in summer 2023 and both were visually assessed. Metabolomics analysis and microscopy observations were completed for one bioassay.

### Metabolomics analysis

The quantification of *P. viticola* biomarkers was based on ultra-high performance liquid chromatography (LC) coupled with mass spectrometry (MS), as previously described by Negrel et al. [18], with some modifications. Metabolites were extracted from the freeze-dried and ground leaf discs (lyophilised for 24 h and processed twice at 30 Hz for 30 s with a stainless-steel bead of 3.0 mm in a Tissue-Lyser II instrument - Qiagen, Hilden, Germany) with methanol (MeOH) using 30 μL per mg of dry weight. The suspension of leaf powder and MeOH was sonicated for 15 min in an ultrasound bath and subjected to centrifugation at 13,300 rcf for 10 minutes, from the resulting solution, 50 μL were transferred to vials.

LC separation was performed using a Vanquish Flex binary UHPLC system (Thermo Fisher Scientific, Waltham, MA, USA) on a Nucleodur C18 HTec column (30 × 2mm, 1.8μm particle size; Macherey- Nagel, Düren, Germany) maintained at 40°C and a mobile phase consisted of methanol/water (7/3) with formic acid (0.1%, v/v) at a flow rate of 0.60 mL/min. The gradient elution programme was as follows: 0 to 0.7 min, 100% A; 0.7 to 2.5 min, 0% A; 2.5 to 3.5 min, 0% A isocratic; 3.5 to 4.5 min, 100% A. he injected volume was 1 µl. The LC system was coupled to an Exploris 120 Q-orbitrap (Thermo Fisher Scientific, Waltham, MA, USA) equipped with an atmospheric pressure chemical ionization (APCI) source operating in positive mode. Parameters were set at 300°C for ion transfer capillary temperature and the corona discharge current was set at 4 μA. Nebulization with nitrogen sheath gas and auxiliary gas were maintained at 30 and 10 arbitrary units, respectively, and the nebulizer temperature was maintained at 400°C. The spectra were acquired within the m/z mass range of 90–1,200 atomic mass units (u.), using a resolution of 60,000 at m/z 200 amu. The system was calibrated internally using EASY-IC calibrating source allowing single mass calibration for full mass range, giving a mass accuracy lower than 1 ppm.

The targeted metabolomics analysis of 12 *P. viticola* biomarkers (Supplementary File 5a), including derivatives of eicosapentaenoic (EPA) and arachidonic acids (AA) and ceramides (Cer), was completed using Xcalibur 4.5 software (Thermo Fisher Scientific, Waltham, MA, USA). The integration of each peak was manually checked before validation. Not found-missing data for targeted metabolomics analysis were imputed with the methodology described by Wei et al. [54], and data were log10 transformed to better satisfy the assumptions of the statistical analysis.

### Statistical analysis

Statistical analysis were carried out using R software [55]. *P. viticola* metabolic biomarker quantification for single leaf discs was tested for Pearson correlation with the OIV452-1 scores excluding “not found” peaks and areas data below 10,000 AU (Supplementary File 5c-5d). The OIV452-1 scores and the *P. viticola* biomarker Cer(d16:1/22:0) data were used in independent t-tests (with Welch’s approximation) to compare *P. viticola* strains infection on the different *Rpv* combinations (Supplementary File 4b-8a). The investigation on the *Rpv* contribution to the resistance to *P. viticola* was carried out with strain-independent ANOVA for the OIV452-1 scores and the *P. viticola* biomarker Cer(d16:1/22:0), considering *Rpv1*, *Rpv3.1* and *Rpv10* (two levels with complete cross interactions), and the Experiment for visual data, as factors (Supplementary File 4c-8b). Homoscedasticity and normality of the residuals were visually checked. Estimated marginal means and standard errors of the *Rpv* combinations for the response variables were calculated by using the ‘emmeans’ package [56] and used to compare the *Rpv* combinations with Tukey’s tests (p-value with the Benjamini-Hochberg correction; Supplementary File 4d; 8c). All data analysed in the manuscript are available in the Supplementary File 9.

## Acknowledgements

The authors thank Claire Villeroy for the help in processing the samples for metabolomic analysis (INRAE UMR1131 SVQV, Colmar, France); the staff of Unité Expérimentale Agronomique et Viticole (UE0871, INRAE-Centre Grand Est-Colmar, France) for maintenance of the plant material; and the VEGOIA phenotyping platform (INRAE-Centre Grand Est-Colmar, France), part of the Strasbourg University ‘Cortecs’ network (https://cortecs.unistra.fr).

## Authors’ contributions

PH, DM, SW and JB conceived the research; PH, DM, JB and RF acquired the financial support; RB, SW and PH supervised the research work; ED, PM and DM conceived the generation of the offspring; RF isolated the *P. viticola* strains virulent on *Rpv10* (*avrRpv10-*); TP, SW, MCL, MAD and MB performed the phenotyping; TP and RB performed the metabolomic analyses; TP, MB and EG were involved in data analysis; TP produced the statistical analysis and the visualizations; TP, SW, RB, PH, ED, PM, JB, MB and EG were involved in data and results interpretation; TP drafted the manuscript with inputs from all authors; all the authors reviewed, edited and approved the final version of the manuscript.

## Data availability statement

All the data are included in the article and its supplementary files.

## Supplementary Files

**Supplementary File S1. Overview of the study and its experimental and technical design.** Summary of the information on the plant material, the pathogen source, the phenotyping strategy and the studied variables of the research. A total of 3 genotypes for each possible combination of the studied resistance loci to *Plasmopara viticola* (*Rpv*) *Rpv1*, *Rpv3.1* and *Rpv10* were challenged with a naive *P. viticola* strain, or avirulent on all the studied *Rpv*, and a *P. viticola* strain virulent toward *Rpv10* (*avrRpv10-*). A total of 3 discs/genotype were visually assessed from 3 to 6 days post inoculation (dpi) recording the incidence and severity of *P. viticola* sporulation, according to the OIV452-1 descriptor, and the incidence of plant necrosis. A total of 3 leaf discs/genotype/strain/time and 2 mock (water)-inoculated discs/genotype/time were collected at 6-12 hours post inoculation (hpi) and at 1-2-3-6 dpi for metabolomics-based phenotyping (discs collected at 6 dpi were those visually assessed). For metabolomics, a methanol (MeOH)-based extraction protocol and high-resolution liquid chromatography-based mass spectrometry (LC-MS) with an ‘Atmospheric pressure chemical ionization’ (APCI) source were adopted for the analysis, and a targeted quantification was performed for 12 *P. viticola*-specific metabolic biomarkers.

**Supplementary File S2. Progression of *Plasmopara viticola* infection on grapevine leaf discs.** Representative disc between 3 and 6 day post inoculation (dpi) for non-resistant (*Rpv-*) and single *Rpv* combinations (*Rpv1*, *Rpv3.1* and *Rpv10*) inoculated with the two *P. viticola* strains of the study (naive and *avrRpv10-*). The discs evidence a different severity and incidence of *P. viticola* sporulation and plant necrosis depending on the *Rpv*-*P. viticola* interaction considered.

**Supplementary File S3. OIV452-1 scores progression on *Plasmopara viticola*-infected grapevine leaf discs.** OIV452-1 mean scores (points) and confidence limits (dashed area; 95%) at 3-4-5-6 days post inoculation (dpi) for each *Rpv* combination - *P. viticola* strain (naive and *avrRpv10-*) interaction studied (1 = sporulation covering the whole disc area; 3 = sporulation covering the most of the disc area in large patches; 5 = sporulation present in delimited patches; 7 = sparse sporulation; and 9 = no sporulation). A total of 3 genotypes/*Rpv* combination and 3 discs/genotype/dpi were assessed in two experiments (Exp; different colors) by stereomicroscope observations.

Supplementary File S4a. Data for the variables characterizing the outcomes of the *Rpv combination*-*Plasmopara viticola* strain interactions. Mean value for the OIV452-1 descriptor, *P. viticola* sporulation incidence (Spo_inc; i.e., OIV452-1 ≠ 9) and plant necrosis incidence (Nec_inc) at 6 days post inoculation (dpi), and sporulation latency period (dpi), for the different *Rpv* combination - *P. viticola* strain interactions studied and for the two experiments carried out (Exp1 and Exp2).

**Supplementary File S4b. T-test comparison of *Plasmopara viticola* strains based on OIV452-1 scores.** T-test (with Welch approximation) to compare *P. viticola* strains (naive and *avrRpv10-*) severity of infection were carried out by using the OIV452-1 scores collected at 6 days post inoculation (dpi) on the different *Rpv* combinations in two experiments (3 genotypes/*Rpv* combination and 3 leaf discs/genotype/strain/experiment). The plots present the OIV452-1 data distribution (box-plot), the mean and standard error of the mean (grey squares and error bars) for the *Rpv* combinations under infection with the two *P. viticola* strains (naive = red; *avrRpv10-* = blue).

**Supplementary File S4c. ANOVA results for the OIV452-1 scores.** Detailed results from ANOVA conducted with the OIV452-1 scores at 6 days post inoculation (dpi), and including *Rpv1*, *Rpv3.1* and *Rpv10* presence (two levels with complete cross interactions), and the Experiment, as factors (degree of freedom-df, f value and p values). Independent ANOVA were conducted for the two *Plasmopara viticola* strains (naive and *avrRpv10-*) and data for 3 genotypes/*Rpv* combination and 3 discs/genotype/strain/dpi of two experiments were used. The plots present the OIV452-1 data distribution (box-plot) and the estimated marginal means and standard errors (black dots and error bars) for the factors investigated in the complete ANOVA analysis for the two *P. viticola* strains (naïve = red; *avrRpv10-* = blue).

**Supplementary File S4d. Pairwise comparisons of *Rpv* combinations based on OIV452-1 scores.** Pairwise comparisons (PWC) between *Rpv* combinations carried out after *Plasmopara viticola* strain (naive and *avrRpv10-*)-independent ANOVA analysis of OIV452-1 scores collected at 6 days post inoculation (dpi) by using the estimated marginal means and standard errors (Tukey’ test with p values with Benjamini-Hochberg correction). The plots present the OIV452-1 data distribution (box-plot) and the estimated marginal means and standard errors (black dots and error bars) for the *Rpv* combinations under infection with the two *P. viticola* strains (naive = red; *avrRpv10-* = blue).

**Supplementary File S5a. Complete list of *Plasmopara viticola*-specific metabolic biomarkers ions quantified by targeted metabolomics analysis.**

**Supplementary File S5b. *Plasmopara viticola* metabolic biomarkers detection.** Percentage (%) of “not found” peaks, and consequent missing area values, for the main different studied conditions: the eight different *Rpv* combinations infected with the two *P. viticola* strains at 3 and 6 days post inoculation (dpi). A total of 3 genotypes/*Rpv* combination, 3 leaf discs/genotype/strain/time and 2 mock (water)-inoculated discs/genotype/time for one experiment were analyzed.

**Supplementary File S5c. Correlations between the OIV452-1 scores and *Plasmopara viticola* metabolic biomarkers abundances.** a) Correlation coefficients (Pearson correlation) between the visual variable OIV452-1 and *P. viticola*-specific metabolic biomarkers for seven retained biomarkers (peak area data were log10-transformed) at 6 days post inoculation (dpi) in 72 samples. b) *P. viticola* strain (naive = red; *avrRpv10-* = blue)-specific correlations between the OIV452-1 scores the metabolic biomarkers abundances.

**Supplementary File S5d. Correlations between *Plasmopara viticola* metabolic biomarkers**. Correlation plots, coefficients and significances (Pearson correlation, p-value < 0.05 for *; < 0.01 for ** and < 0.001 for ***) between the six most informative *P. viticola*-specific metabolic biomarkers (peak area data log10-transformed) at 6 days post inoculation (72 samples).

**Supplementary File 6. *Plasmopara viticola* metabolic biomarkers abundances progression in grapevine leaf discs.** The heatmap shows the mean peak area (log10 of the arbitrary unit) for the six most informative *P. viticola*-specific metabolic biomarkers (Eicosapentaenoic acid - EPA, Eicosapentaenoyl-glycerol - EPG, Arachidonic acid - AA and the three ceramides Cer(d16:1/18:0), Cer(d16:1/18:0) and Cer(d16:1/18:0); right labels) quantified in the discs of the different *Rpv* combination (bottom labels) - *P. viticola* strain (left labels) interactions studied at 6-12 hours post inoculation (hpi; top labels) and at 0-1-2-3-6 days post inoculation (dpi; top labels). A total of 9 discs/*Rpv* combination/strain/time for the inoculated samples, and 6 discs/*Rpv* combination/time for the mock (water)-inoculated samples were analyzed by high-resolution liquid chromatography-based mass spectrometry (LC-MS).

**Supplementary File S7a. *Plasmopara viticola* metabolic biomarkers abundances ratios comparing *P. viticola* strains.** Mean peak area data per *Rpv* combination per *P. viticola* strain (naive and *avrRpv10-*) at 3 and 6 days post inoculation (dpi) for the six *P. viticola* metabolic biomarkers analyzed and log2 ratios between the mean metabolite peak areas for the same *Rpv* combination (3 genotypes/*Rpv* and 3 discs/genotype/strain/time; table and heatmap). The ratios highlight the different aggressiveness of the *P. viticola* strains on the studied *Rpv* combinations.

**Supplementary File S7b. *Plasmopara viticola* metabolic biomarkers abundances ratios comparing the infection on the *Rpv* combinations.** Mean peak area data per *Rpv* combination per *P. viticola* strain (naive and *avrRpv10-*) at 3 and 6 days post inoculation (dpi) for the six *P. viticola* metabolic biomarkers analyzed and log2 ratios between the mean areas for different *Rpv* combinations (3 genotypes/*Rpv* and 3 discs/genotype/strain/time; table and heatmap). The ratios highlight the different efficacy of *Rpv* combinations in limiting the infection of the *P. viticola* strains.

**Supplementary File S8a. T-test comparison of *Plasmopara viticola* strains based on the abundances of *P. viticola* metabolic biomarkers.** T-tests (with Welch approximation) to compare *P. viticola* strains (naive and *avrRpv10-*) severity of infection were carried by using the peak area data (log10 of the arbitrary unit-AU) for the most informative *P. viticola* metabolic biomarker Cer(d16:1/22:0) collected at 3 and 6 days post inoculation (dpi) on the different *Rpv* combinations (3 genotypes/*Rpv* combination and 3 leaf discs/genotype/strain). The plots present the Cer(d16:1/22:0) data distribution (box-plot), the mean and standard error of the mean (grey squares and error bars) for the *Rpv* combinations under infection with the two *P. viticola* strains (naive = red; *avrRpv10-* = blue).

**Supplementary File S8b. ANOVA results for the *Plasmopara viticola* metabolic biomarkers.** Detailed results from ANOVA conducted with the most informative *P. viticola*-metabolic biomarker Cer(d16:1/22:0) abundances (log10 of the arbitrary unit-AU) at 3-6 days post inoculation (dpi), and including *Rpv1*, *Rpv3.1* and *Rpv10* presence (two levels with complete cross interactions), and the Experiment, as factors (degree of freedom-df, f value and p values). Independent ANOVA were conducted for the two *P. viticola* strains (naive and *avrRpv10-*) and data for 3 genotypes/*Rpv* combination and 3 discs /genotype/strain/dpi of one experiment were used. The plots present the Cer(d16:1/22:0) abundances distribution (box-plot) and the estimated marginal means and standard errors (black dots and error bars) for the factors investigated in the complete ANOVA analysis for the two *P. viticola* strains (naive = red; *avrRpv10-* = blue).

**Supplementary File S8c. Pairwise comparisons between Rpv combinations for the *Plasmopara viticola* Cer(d16:1/22:0) biomarker.** Pairwise comparisons (PWC) between *Rpv* combinations carried out after *P. viticola* strain (naive and *avrRpv10-*)-independent ANOVA analysis of Cer(d16:1/22:0) peak area data (log10 of the arbitrary unit; log10 AU) collected at 3 and 6 days post inoculation (dpi) by using estimated marginal means and standard errors (Tukey’ test with p-values with Benjamini-Hochberg correction). The plots present the Cer(d16:1/22:0) data distribution (box-plot) and the estimated marginal means and standard errors (black dots and error bars) for the *Rpv* combinations under infection with the two *P. viticola* strains (naive = red; *avrRpv10-* = blue).

**Supplementary File S9a. Data for visually-assessed variables.** Scores for single leaf disc (DiscID) for the variables OIV452-1 (1=extend and dense sporulation/high susceptibility; 9=absence of sporulation/total resistance) and necrosis incidence (0=absence; 1=presence) collected at different days post inoculation (dpi).

**Supplementary File S9b. Data for *Plasmopara viticola* metabolic biomarkers quantification.** Peak area data, or abundances, (arbitrary unit-AU) with missing values (not found peaks) attributed as described in Wei et al. (2018) for the *P. viticola* metabolic biomarkers analyzed: eicosapentaenoic acid (EPA), eicosapentaenoyl-glycerol (EPG=, dieicosapentaenoyl-glycerol (DEPG), trieicosapentaenoyl-glycerol (TEPG), arachidonic acid (AA), arachidonoyl-glycerol (AG), diarachidonoyl-glycerol (DAG), triarachidonoyl-glycerol (TAG), and ceramides Cer(d16:1/16:0), Cer(d16:1/18:0), Cer(d16:1/20:0) and Cer(d16:1/22:0). Sample data correspond to single leaf discs collected at different hours or days post the inoculation (hpi and dpi) with a *P. viticola* strain (naive or *avrRpv10-*) or water (mock).

**Supplementary File S9c. Data to study the correlation between the OIV452-1 scores and the abundances of *Plasmopara viticola* metabolic biomarkers.** Data for the OIV452-1 descriptor and the most informative *P. viticola* biomarkers for the leaf discs/samples collected at 6 dpi and used to investigate the correlation between variables used to assessed the severity of *P. viticola* infection.

