## Supplementary material for "Metabolic biomarker-based phenotyping unveils quantitative effects of plant resistance and pathogen aggressiveness in the grapevine (Vitis spp.) - downy mildew (Plasmopara viticola) pathosystem": Supplementary File S3.pdf

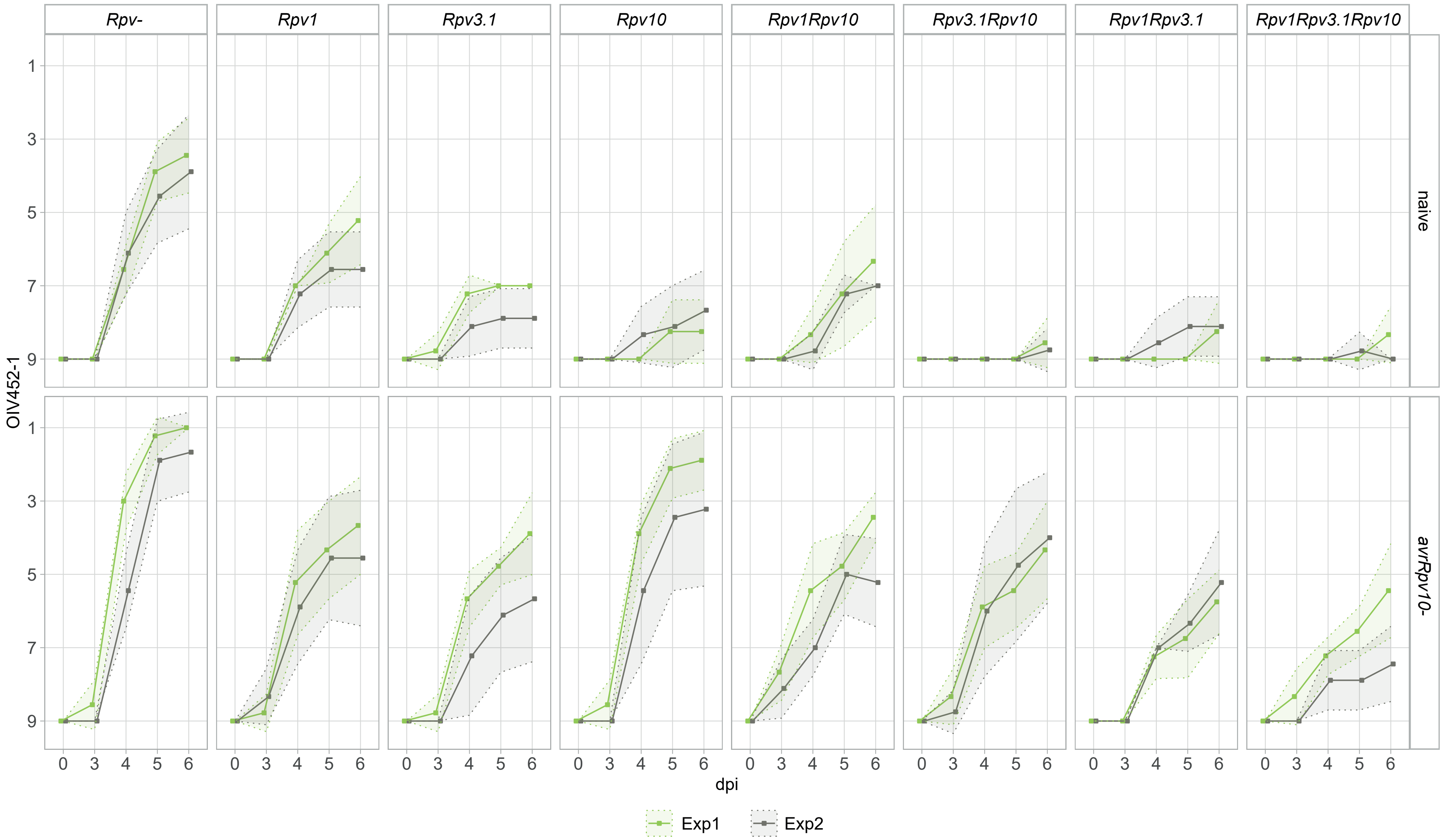


**Supplementary File S3. OIV452-1 scores progression on *Plasmopara viticola*-infected grapevine leaf discs.** OIV452-1 mean scores (points) and confidence limits (dashed area; 95%) at 3-4-5-6 days post inoculation (dpi) for each *Rpv* combination - *P. viticola* strain (naive and *avrRpv10-*) interaction studied (1 = sporulation covering the whole disc area; 3 = sporulation covering the most of the disc area in large patches; 5 = sporulation present in delimited patches; 7 = sparse sporulation; and 9 = no sporulation). A total of 3 genotypes/*Rpv* combination and 3 discs/genotype/dpi were assessed in two experiments (Exp; different colors) by stereomicroscope observations.
