## Supplementary material for "Metabolic biomarker-based phenotyping unveils quantitative effects of plant resistance and pathogen aggressiveness in the grapevine (Vitis spp.) - downy mildew (Plasmopara viticola) pathosystem": Supplementary File S6.pdf

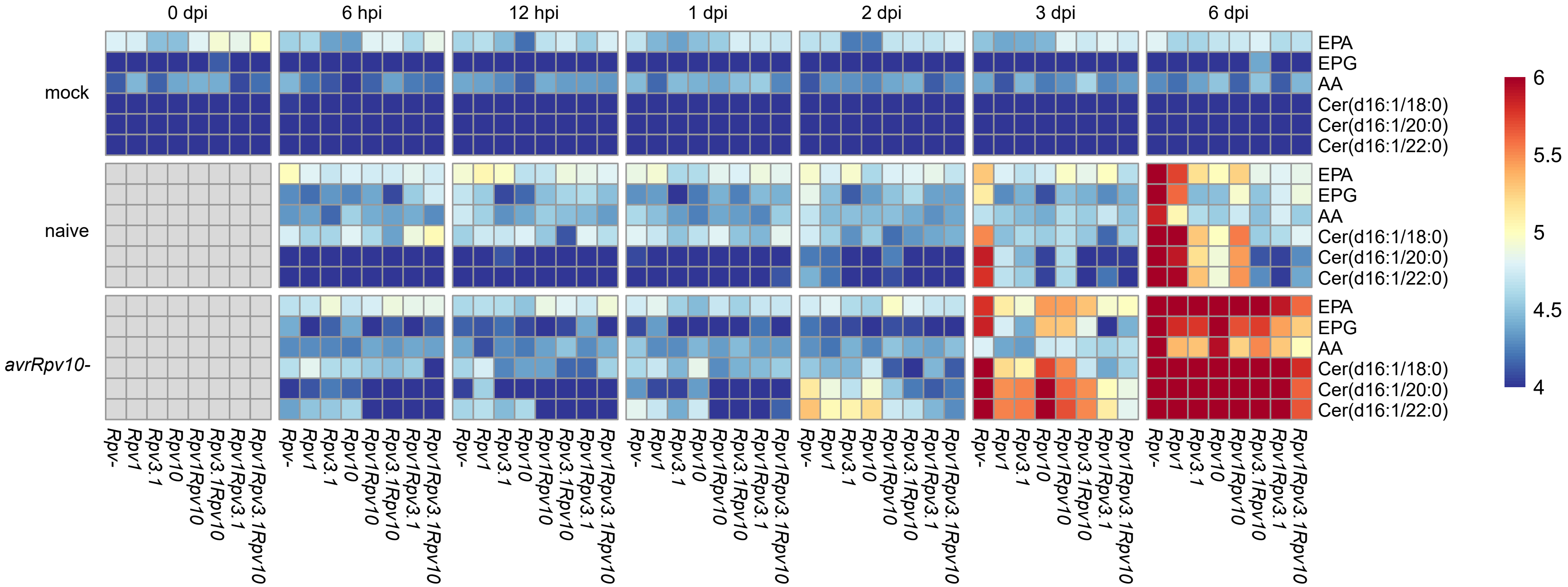


**Supplementary File 6. *Plasmopara viticola* metabolic biomarkers abundances progression in grapevine leaf discs.** The heatmap shows the mean peak area (log10 of the arbitrary unit) for the six most informative *P. viticola*-specific metabolic biomarkers (Eicosapentaenoic acid - EPA, Eicosapentaenoyl-glycerol - EPG, Arachidonic acid - AA and the three ceramides Cer(d16:1/18:0), Cer(d16:1/18:0) and Cer(d16:1/18:0); right labels) quantified in the discs of the different *Rpv* combination (bottom labels) - *P. viticola* strain (left labels) interactions studied at 6-12 hours post inoculation (hpi; top labels) and at 0-1-2-3-6 days post inoculation (dpi; top labels). A total of 9 discs/*Rpv* combination/strain/time for the inoculated samples, and 6 discs/*Rpv* combination/time for the mock (water)-inoculated samples were analyzed by high-resolution liquid chromatography-based mass spectrometry (LC-MS).
