## Supplementary material for "Metabolic biomarker-based phenotyping unveils quantitative effects of plant resistance and pathogen aggressiveness in the grapevine (Vitis spp.) - downy mildew (Plasmopara viticola) pathosystem": Supplementary Files Index.pdf

Supplementary File S1.
Overview of the study and its experimental and technical design.
Supplementary File S2.
Progression of *Plasmopara viticola* infection on grapevine leaf discs.
Supplementary File S3.
OIV452-1 scores progression on *Plasmopara viticola*-infected grapevine leaf discs.
Supplementary File S4
Supplementary File S4a. Data for the variables characterizing the outcomes of the *Rpv combination-Plasmopara viticola* strain interactions.
Supplementary File S4b. T-test comparison of *Plasmopara viticola* strains based on OIV452-1 scores.
Supplementary File S4c. ANOVA results for the OIV452-1 scores.

Supplementary File S4d. Pairwise comparisons of *Rpv* combinations based on OIV452-1 scores.

Supplementary File S5

Supplementary File S5a. Complete list of *Plasmopara viticola*-specific metabolic biomarkers ions quantified by targeted metabolomics analysis.

Supplementary File S5b. *Plasmopara viticola* metabolic biomarkers detection.

Supplementary File S5c. Correlations between the OIV452-1 scores and *Plasmopara viticola* metabolic biomarkers abundances.

Supplementary File S5d. Correlations between *Plasmopara viticola* metabolic biomarkers.

Supplementary File 6.

*Plasmopara viticola* metabolic biomarkers abundances progression in grapevine leaf discs.

Supplementary File 7

Supplementary File S7a. *Plasmopara viticola* metabolic biomarkers abundances ratios comparing *P. viticola* strains.

Supplementary File S7b. *Plasmopara viticola* metabolic biomarkers abundances ratios comparing the infection on the *Rpv* combinations.

Supplementary File 8

Supplementary File S8a. T-test comparison of *Plasmopara viticola* strains based on the abundances of *P. viticola* metabolic biomarkers.

Supplementary File S8b. ANOVA results for the *Plasmopara viticola* metabolic biomarkers.

Supplementary File S8c. Pairwise comparisons between Rpv combinations for the *Plasmopara viticola* Cer(d16:1/22:0) biomarker.

Supplementary File 9

Supplementary File S9a. Data for visually-assessed variables.

Supplementary File S9b. Data for *Plasmopara viticola* metabolic biomarkers quantification.

Supplementary File S9c. Data to study the correlation between the OIV452-1 scores and the abundances of *Plasmopara viticola* metabolic biomarkers.
