## Supplementary figures and images for "Metabolic biomarker-based phenotyping unveils quantitative effects of plant resistance and pathogen aggressiveness in the grapevine (Vitis spp.) - downy mildew (Plasmopara viticola) pathosystem"

### Supplementary File S1.jpg

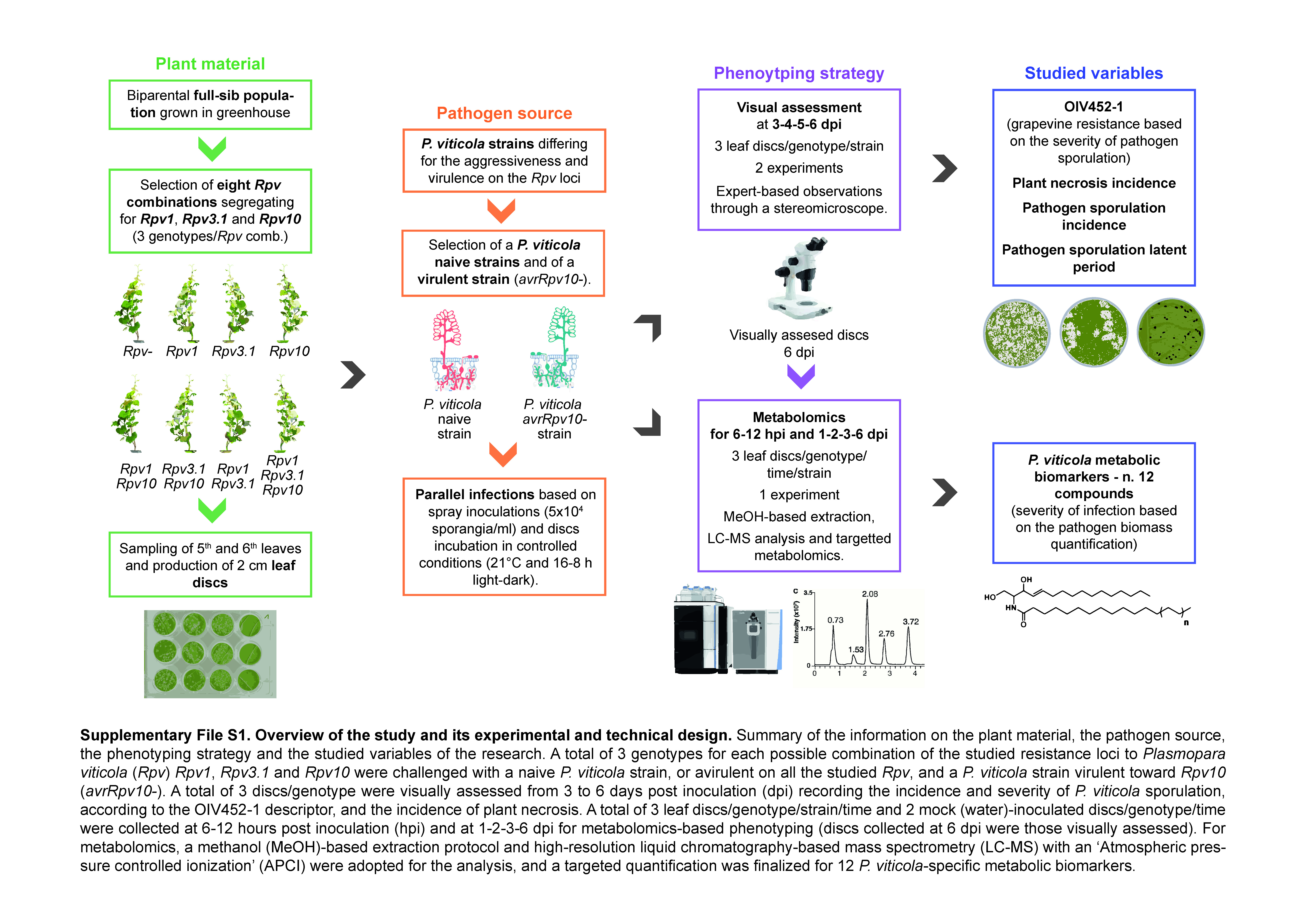

### Supplementary File S2.jpg

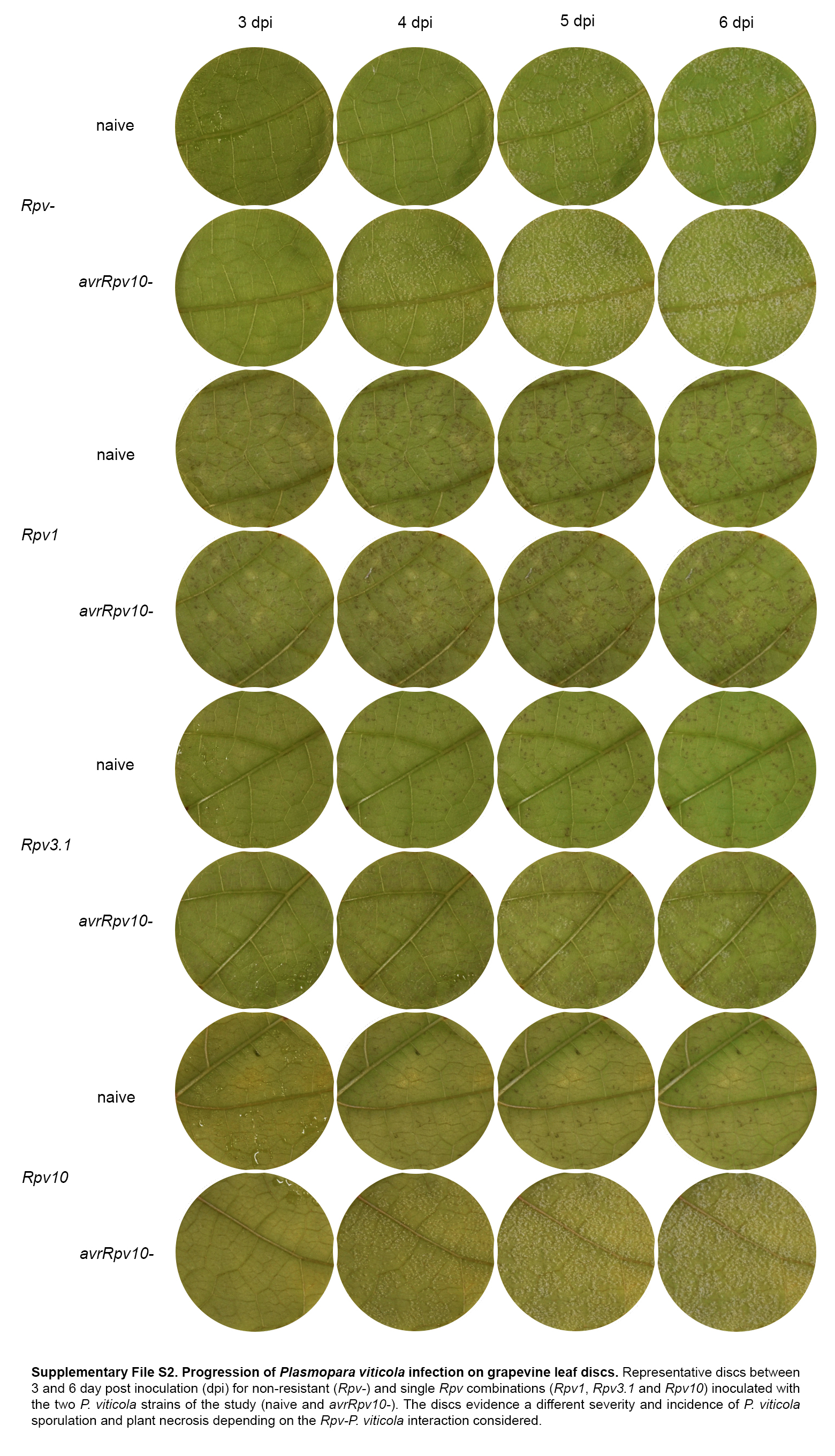
